# Bioactivity of Areca catechu and Azadirachta indica seed oil against Bulinus sp, an intermediate host of Schistosoma haematobium

**DOI:** 10.1101/2023.10.20.563218

**Authors:** Katamssadan Haman Tofel, Gillian Awafor, Joan Echi Eyong Ebanga, Ngum Helen Ntonifor

**Author notes:** These authors contributed equally to this work. These authors also contributed equally to this work.

## Abstract

**Background:** Schistosomiasis is a parasitic disease transmitted by freshwater snail with infected *Schistosoma* parasites. The disease is endemic in many parts of Asia, Africa, and South America affecting people who are unable to avoid contact with water, either because of their profession or because of lack of reliable source of safe water for drinking, washing and bathing.

**Methods:** Seed oil extract of *Azadirachta indica* and the aqueous and ethanol extracts of *Areca catechu* were tested on freshwater snails *Bulinus* sp. Bioassays were performed on adults and eggs of snails at varying concentrations (50, 100, 200, 400, 800 and 1600mg/L for the extracts of *A. catechu* and 1, 2.5, 5, 7 and 10mL/L for *A. indica* extract).

**Results:** The result revealed that mortality increased with ascending concentration. There was significant difference in the molluscicidal activities of the three extracts (*p* < 0.05). Among the tested the plant extracts, *A. indica* seed oil showed the highest molluscicidal activity (100%) from 7 mL/L. Meanwhile, the lowest molluscicidal activity (2.5%) was found in the *A. catechu* aqueous extract. It was also observed that the potency of the extracts to inhibit egg hatching increases as the concentrations increased.

**Conclusion:** *Azadirachta indica* seed oil and the ethanolic extract of *A. cathetu* could be considered as a veritable means of controlling schistosomiasis vector. Also, in endemic areas where communities are likely to accept the use of local plants, the studied plants stand as good candidates to replace the conventional medicines which are expensive.

**Synopsis:** 

## Introduction

Schistosomiasis is caused by schistosomes which belong to the family Schistosomatidae that, includes species that are among the most dreaded parasites of humans [1]. Five clinically important species cause majority of human infections [2]. *Schistosoma m*ansoni, *Schistosoma japonicum, Schistosoma mekon*gi *Schistosoma intercalatum* and *Schistosoma haematobium*. Schistosomiasis is considered as neglected tropical disease and affects more than 250 million people in tropical and sub-tropical regions of the world, in poor communities without potable water or adequate sanitation. Sub-Saharan Africa accounts for approximately 90% of worldwide cases [3]. Disease assessments indicate that schistosomiasis accounts for up to seventy million disability adjusted life years lost annually, considering the amount of end organ pathologies in the liver for *S. mansoni* and *S. japonicum* and the bladder and kidney for *S. haematobium* coupled with chronic morbidities associated with impaired child growth and development, chronic inflammation, anaemia and other nutritional deficiencies [4]. The freshwater planorbid snails; *Bulinus* sp and *Biomphalaria* sp are the intermediate hosts of *S. haematobium* and *S. mansoni*, which cause urinary and intestinal schistosomiasis respectively in Cameroon.

Snail control is one of the methods of choice for the fight against schistosomiasis transmission and might include the use of chemical molluscicides, bio-agents and environmental management. Chemical control depends on the elimination of snails from the habitat by molluscicides. Different molluscicides have been tested in the laboratory and in the field. They are either synthetically manufactured or of plant origin. The synthetic molluscicide, niclosamide (Bayluscide®), is one of the most used chemicals because it fulfils most of the characteristics of the ideal molluscicide. Although it does not harm crop plants, it is slowly absorbed through the intestine, skin and mucous membranes of mammals. Amphibians and fish are, however, very sensitive to it because the chemical has a strong irritant effect on their mucosal membranes [5]. Thus, an alternative control tool is desirable also because niclosamide has to be imported and it is expensive and has an environmental impact, even if small. Furthermore, applying only a single molluscicide such as niclosamide could possibly result in snails developing resistance to it [6].

Recently, molluscicidal plants have been drawing increasing attention for their environmental friendliness, accessibility, and cost effectiveness. They are particularly suitable for community-based snail control activities in places where schistosomiasis transmission is more focal [7]. Snail control by using synthetic molluscicides also plays an important role in the integrated control programme for schistosomiasis. Much attention has been given to the study of plant molluscicides because they may provide a cheap, biodegradable and effective control approach in rural areas of developing countries, where schistosomiasis is endemic. However, the toxicity of these molluscicides to non-target organisms and ecosystem destruction may render them less efficient. Commercial molluscicides are not largely used as major control mechanisms due to their cost implications and toxicity effects on non-target organisms such as fish. Therefore, there is a need to search for cheap and safe molluscicides [8]. This is the basis on which the experiment was conducted using extracts of *Azadirachta indica* and *Areca catechu*. In addition, the control of the intermediate host in the control of schistosomiasis has been shown to be of paramount importance to prevent re-infection after treatment of humans.

## Materials and Methods

### Collection and preparation of plant extracts

Mature seeds of *A. catechu* were locally collected based on ethnobotanical information. The plant parts were collected around the University of Bamenda campus (N 5?59’22” and E 10?152”)? and taken to the laboratory. The identified plant seeds were air dried and ground into fine powder using an electric blender, the powder was weighed and collected into clean cellophane bags, labeled and kept in a cool dry place until further use. Neem seed oil was obtained from Maroua, Far North region of Cameroon.

Four hundred grammes of *A. catechu* powder was soaked in 2 L of 70% ethanol for two days (48 h), with constant shaking. The solution was filtered using filter paper (Whatman No. 1) and dried in laboratory oven at 60ºC. The dried materials constituted the ethanol extract [9]. Similarly, 400 g of *A. catechu* powder was soaked in 2 L of distilled water for two days (48 h) polar compounds extraction. The solution was filtered using filter paper (Whatman N°1) and dried. The dried material constituted the aqueous extract [9].

### Collection, sampling and breeding of snails

The snails used in this study were *Bulinus* sp. Six hundred adult snails of the genus *Bulinus* were collected from their natural habitat in Bambui stream around the school of Agriculture of the University of Bamenda, kept in a clean aquarium and taken to the laboratory. In the laboratory, the snails were identified using the snail identification key of Danish Bilharzias Laboratory [10]. In the laboratory, the snails were acclimatized for 72 h with dechlorinated tap water and maintained in aquaria of plastic bowls (12cm depth×30cm diameter with a capacity of about 5 liters). Snails were fed on fresh lettuce after removing their midrib [11]. The aquaria were maintained with constant light for 12 h daily (12 h light and 12 h dark). Water was changed once a week or when necessary. Dead snails were removed as soon as possible from the troughs to prevent water fouling.

### Preparation of egg-masses

Ten adult *Bulinus* snails taken from the stock aquaria were transferred to plastic trays (12 cm depth×30 cm diameter) containing two liters of water. Then, 2-3 pieces of polythene sheets (about 5cm× 15cm) were place in each plastic tray. The snails were fed with fresh lettuce and allowed to lay eggs. The polythene sheets were checked for egg masses. After this period the snails were transferred to a new plastic tray and the same steps were repeated. Polythene sheet containing the egg-masses were easily located and isolated by cutting the plastic around each egg-mass with a scalpel (about 0.5-1.0cm from the egg-mass). The egg-masses, each attached to a piece of polythene, was immersed three times in different petri-dishes containing clean water to remove any debris and transferred to containers containing 200 mL of dechlorinated tap water and the dishes were covered.

### Bioassays

Molluscicidal evaluation was performed according to the WHO recommended guidelines for molluscicidal tests [12]. Batches of 20 snails were pooled together in each clean 200 mL plastic containers. The setups were left for 24 h and the snails were fed on fresh lettuce. Different concentrations of 50, 100, 200, 400, 800 and 1600 mg/L of both the aqueous and the ethanol extracts of *A. catechu* and 1, 2.5, 5, 7 and 10 mL/L *of A. indica* oil extracts were made. Tween 20 was used to mix water with neem seed oil. A control was setup with 100 mL of water only. After 24 h, the distilled water was discarded from the containers holding 20 snails each and replaced with 200 mL for the different extract’ concentrations, and the control preparation. Four replicates were set for each of the different concentrations made. The setups were left for 24 h and after this time the plant extracts were replaced with 100 mL water and mortality recorded after additional 24 h. This was done by pocking the foot of the snail with a wooden applicator stick where lack of motion signified death of the snail [13]. The number of dead or surviving snails was recorded for each of the treatments and controls. The egg masses were cleaned and placed in groups of 10 in different plastic containers holding 100 mL of distilled water. The same procedure used for adult snails was applied and the egg masses observed for hatching. Failure of egg masses to hatch within ten days signified mortality.

### Statistical Analysis

Data on % corrected mortality and % egg inhibition were arcsine [(square root(x/100)] transformed and the number of eggs hatched were log (x + 1) transformed to homogenize the variance. The transformed data were subjected to the ANOVA procedure using the Statistical Analysis System (SAS). Tukey (HSD) test (P = 0.05) was applied for mean separation. Probit analysis was applied to determine the lethal concentrations causing 50% (LC50) and 95% (LC95) mortality of snails. Abbott’s formula was used to correct for control mortality before Probit analysis and ANOVA.

## Results

### Mortality of adult snails

Tables 1, 2 and 3 below shows the average mortality of the snails subjected to different concentrations of the various concentrations of aqueous and ethanol extracts of A. catechu and seed oil of *A. indica* respectively. The result revealed that mortality increased with increased concentration. Aqueous extracts of *A. catechu* recorded an average percentage mortality of 37.5%, 35% and 70% for 400, 800 and 1600 mg/L respectively. Similarly with the ethanol extract of *A. catechu* it was observed that none of the snail died when they were exposed to 50 mg/L of the extract. Meanwhile, as the concentrations increased from 100 mg/L to 400 mg/L, mortality also increased from 12.5% to 17.5%. For 800 mg/L, 35% mortality was recorded, the highest mortality was observed with 1600 mg/L (58.33%). *Azadirachta indica* extract exhibited varied degrees of mortality at different concentrations. 23.75% and 27.5% snail mortality were recorded at 1 mL/L and 2.5 mL/L respectively. Similarly, 85% mortality was recorded at 5mL/L. Meanwhile, 100% mortality was only achieved at 10mL/L and 7 mL/L.

**Table 1.** Molluscicidal activity of *A. catechu* aqueous extract on *Bulinus* sp adult snails.

| Concentration mg/L | %Mortality (means ± SE)<sup>#</sup> |
| --- | --- |
| 0 | 0.00 ± 0.00<sup>d</sup> |
| 50 | 2.50 ± 1.44<sup>cd</sup> |
| 100 | 3.75 ± 2.39<sup>cd</sup> |
| 200 | 6.25 ± 2.39<sup>cd</sup> |
| 400 | 37.50 ± 3.54<sup>b</sup> |
| 800 | 35.00 ± 2.89<sup>bc</sup> |
| 1600 | 70.00 ± 2.04<sup>a</sup> |
| F<sub>(6; 21)</sub> | 49.03<sup>***</sup> |

<sup>#</sup>Each value is a mean ± standard error of four replicate. Means followed by the same letter along the column do not differ significantly at $p = 0.05$ (Tukey's test) ns $p > 0.05$ ; \*\*\* $p < 0.001$ .

**Table 2.** Molluscicidal activity of *A. catechu* ethanolic extract on *Bulinus* sp adult snails.

| Concentration mg/L | %Mortality (means ± SE)<sup>#</sup> |
| --- | --- |
| 0 | 0.00 ± 0.00<sup>d</sup> |
| 50 | 0.00 ± 0.00<sup>d</sup> |
| 100 | 12.50 ± 1.44<sup>bc</sup> |
| 200 | 12.50 ± 5.50<sup>bc</sup> |
| 400 | 17.50 ± 1.44<sup>b</sup> |
| 800 | 35.00 ± 2.89<sup>ab</sup> |
| 1600 | 58.33 ± 16.00<sup>a</sup> |
| F<sub>(6; 21)</sub> | 187.10<sup>***</sup> |

<sup>#</sup>Each value is a mean ± standard error of four replicate. Means followed by the same letter along the column do not differ significantly at $p = 0.05$ (Tukey's test) ns $p > 0.05$ ; \*\*\* $p < 0.001$

**Table 3.**
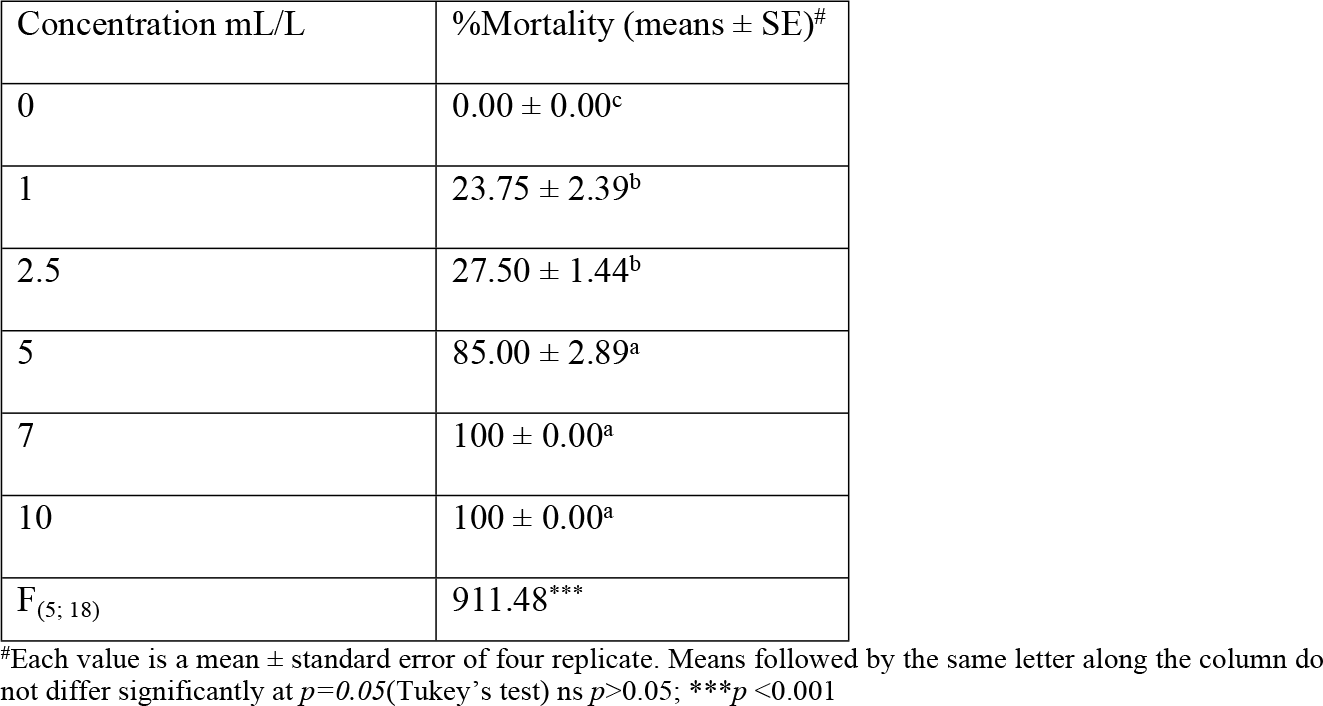
Molluscicidal activity of *A. indica* seed oil on *Bulinus* sp adult snails.

### Mortality of egg masses

Tables 4, 5 and 6 below show the average mortality of the egg masses subjected to different concentrations of the various concentrations of aqueous and ethanol extracts of A. catechu and oil extract of A. indica respectively. Aqueous extracts of A. catechu recorded an average percentage mortality of 76.25%, 88.75% and 98.75% for 50, 100 and 200 mg/L respectively. Similarly, the ethanol extract of A. catechu recorded an average percentage mortality of 67.5%, 82.5% and 95% for 50, 100 and 200 mg/L respectively. In all the tested extracts, the oil extract of A. indica had the highest percentage mortality on the egg masses achieving 91.25% mortality at only 1mL/L.

**Table 4.** Molluscicidal activity of *A. catechu* aqueous extract on *Bulinus sp* egg masses.

| Concentration mg/L | Number of eggs hatched<br>(means ± SE)<sup>#</sup> | % Inhibition of egg development<br>(means ± SE)<sup>#</sup> |
| --- | --- | --- |
| 0 | 20.00 ± 0.00<sup>a</sup> | 0.00 ± 0.00<sup>c</sup> |
| 50 | 23.75 ± 0.25<sup>b</sup> | 76.25 ± 1.25<sup>b</sup> |
| 100 | 11.25 ± 0.25<sup>c</sup> | 88.75 ± 1.25<sup>a</sup> |
| 200 | 1.25 ± 0.25<sup>d</sup> | 98.75 ± 1.25<sup>a</sup> |
| 400 | 0.00 ± 0.00<sup>d</sup> | 100 ± 0.00<sup>a</sup> |
| 800 | 0.00 ± 0.00<sup>d</sup> | 100 ± 0.00<sup>a</sup> |
| 1600 | 0.00 ± 0.00<sup>d</sup> | 100 ± 0.00<sup>a</sup> |
| F<sub>(6; 21)</sub> | 2001.1<sup>***</sup> | 610.42<sup>***</sup> |

<sup>#</sup>Each value is a mean ± standard error of four replicate. Means followed by the same letter along the column do not differ significantly at $p = 0.05$ (Tukey's test) ns $p > 0.05$ ; \*\*\* $p < 0.001$

**Table 5.** Molluscicidal activity of *A. catechu* ethanolic extract on *Bulinus sp* egg masses.

| Concentration mg/L | Number of eggs hatched<br>(means ± SE)<sup>#</sup> | % Inhibition of egg development<br>(means ± SE)<sup>#</sup> |
| --- | --- | --- |
| 0 | 20.00 ± 0.00<sup>a</sup> | 0.00 ± 0.00<sup>c</sup> |
| 50 | 32.50 ± 0.29<sup>b</sup> | 67.50 ± 1.44<sup>b</sup> |
| 100 | 17.50 ± 0.29<sup>c</sup> | 82.50 ± 1.44<sup>a</sup> |
| 200 | 5.00 ± 0.58<sup>d</sup> | 95.00 ± 2.89<sup>a</sup> |
| 400 | 0.00 ± 0.00<sup>d</sup> | 100 ± 0.00<sup>a</sup> |
| 800 | 0.00 ± 0.00<sup>d</sup> | 100 ± 0.00<sup>a</sup> |
| 1600 | 0.00 ± 0.00<sup>d</sup> | 100 ± 0.00<sup>a</sup> |
| F<sub>(6; 21)</sub> | 742.50<sup>***</sup> | 246.72<sup>***</sup> |

<sup>#</sup>Each value is a mean ± standard error of four replicate. Means followed by the same letter along the column are do not differ significantly at $p = 0.05$ (Tukey's test) ns $p > 0.05$ ; \*\*\* $p < 0.001$

**Table 6.**
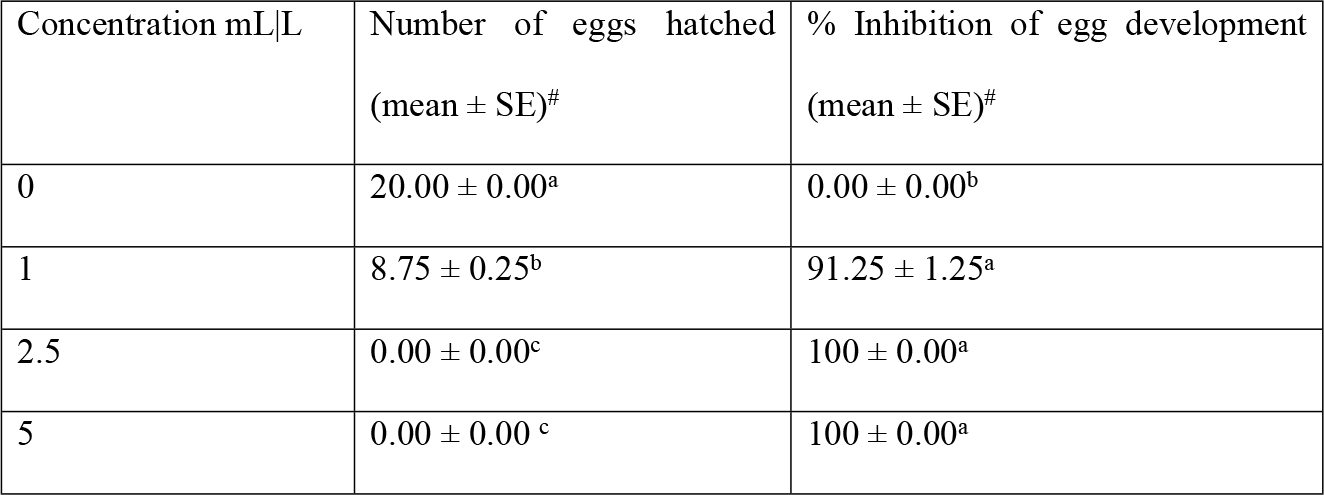
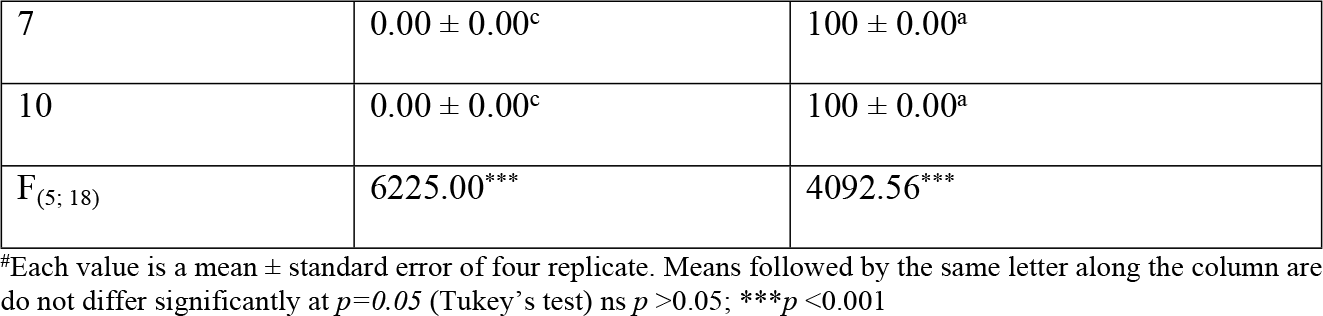
Molluscicidal activity of *A. indica* seed oil on *Bulinus sp* egg masses.

| Concentration mL/L | Number of eggs hatched<br>(mean ± SE)<sup>#</sup> | % Inhibition of egg development<br>(mean ± SE)<sup>#</sup> |
| --- | --- | --- |
| 0 | 20.00 ± 0.00<sup>a</sup> | 0.00 ± 0.00<sup>b</sup> |
| 1 | 8.75 ± 0.25<sup>b</sup> | 91.25 ± 1.25<sup>a</sup> |
| 2.5 | 0.00 ± 0.00<sup>c</sup> | 100 ± 0.00<sup>a</sup> |
| 5 | 0.00 ± 0.00<sup>c</sup> | 100 ± 0.00<sup>a</sup> |
| 7 | 0.00 ± 0.00<sup>c</sup> | 100 ± 0.00<sup>a</sup> |
| 10 | 0.00 ± 0.00<sup>c</sup> | 100 ± 0.00<sup>a</sup> |
| F<sub>(5, 18)</sub> | 6225.00*** | 4092.56*** |

#Each value is a mean ± standard error of four replicate. Means followed by the same letter along the column are do not differ significantly at $p=0.05$ (Tukey's test) ns $p>0.05$ ; \*\*\* $p<0.001$

### Toxicity of plant extracts

The results in table 7 revealed that, the lethal concentration that killed 50% (LC_50_) of the adult B. sp snails, when *A. indica* seed oil, aqueous and ethanol extracts of *A. catechu* was used were: 2.20mL/L, 0.93mg/L and 1.35mg/L, respectively, while the lethal concentration that killed 95% of the snails (LC_95_) were: 7.67mL/L, 7.97mg/L and 21.24mg/L, respectively. While in table 8 it was observed that the lethal concentration that killed 50% (LC_50_) of the egg masses of *Bulinus* sp snails, when *A. indica* extract, aqueous and ethanol extracts of *A. catechu* was used were: 0.80mL/L, 0.03mg/L and 0.03mg/L, respectively, while the lethal concentration that killed 95% of the egg masses (LC_95_) were: 1.04 mL/L, 0.13mg/L and 0.18mg/L, respectively.

**Table 7.** Toxicity of the plant extracts *Bulinus* adult snails.

| Products | Slope ± SE<sup>a</sup> | R<sup>2</sup> | LC<sub>50</sub> (95% FL<sup>b</sup>) | LC<sub>95</sub> (95% FL) | X<sup>2</sup> (p value) |
| --- | --- | --- | --- | --- | --- |
| *A. catechu* aqueous | 1.76 ± 0.30 | 0.77 | 0.93 (0.58 - 2.20) | 7.97 (3.27 - 119.06) | 16.45 (0.02) |
| *A. catechu* ethanol | 1.37 ± 0.25 | 0.78 | 1.35 (0.76 - 5.09) | 21.24 (5.46 - 118.28) | 12.99 (0.01) |
| *A. indica* seed oil | 2.20 ± 0.95 | 0.93 | 2.20 (101.31 - 1545) | 7.67 (3.71 - 137.94) | 56.61(0.01) |

<sup>a</sup>SE= Standard Error; <sup>b</sup>FL = Fiducial limit.

**Table 8.**
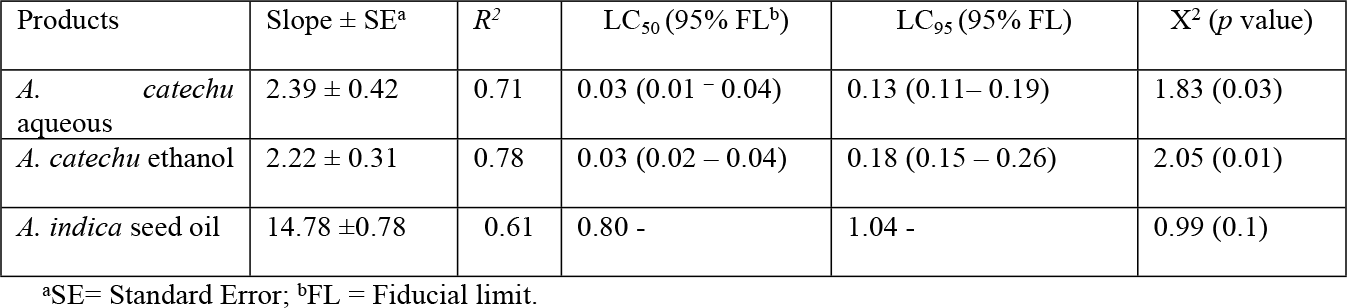
Toxicity of the plant extracts on the egg masses.

## Discussion

The search for bioactive plants components which can be used as non-conventional molluscicides and anti-helminthes has received considerable attention in recent times because of the increasing, worldwide development of resistance to chemical molluscicides in mollusc populations respectively. However, scientific evidence to validate the use of plants remains limited [14]. As the intermediate hosts, molluscs play a major role in the transmission of schistosomes; they are the sites of an intense multiplication of parasites. Thus, snail control strategies are considered a priority for the reduction of schistosomiasis transmission. A standardized procedure devised for the laboratory screening of synthetic chemical molluscicides is used to evaluate plant molluscicides.

This study revealed that the seed oil extract of *A. indica* was highly toxic to freshwater snails that serve as intermediate host of schistosomiasis but, the aqueous and ethanol extracts of *A. catechu* were fairly toxic to the freshwater snails because at the highest concentration only 70% and 58% mortality, respectively, was recorded for all replicates. The toxicity of *A. indica* was most active at the last two concentrations (7 and 10 mL/L) at 24 hours of exposure where 100% mortality was observed. Lower concentrations (1, 2.5 and 5 mL/L) did not give 100% mortality which indicated that the snails were able to withstand the toxicity of the oil extract at lower concentrations. Reduced mortality observed at lower concentrations may be due to low concentration and the time frame, while higher mortality observed at higher concentrations within short time can probably be linked to the oil toxicity and concentration itself.

Plant materials may release active ingredients slowly so that the effect on snail population is delayed, that is, the plant may act as a slow-release matrix. In this mode, there may also be effects on feeding and oviposition. Other plant substances affect orientation and feeding behaviour [15]. A bioassay of whole plants or parts in which snails are killed within 24 hours at a dosage below 100 mg/L indicates that the molluscicide is released quickly and the material may be a good candidate for LC50 determination [16]. This is in agreement with findings of this study in which death of snails was observed within 24 hours exposure to concentrations below 100mg/L of the plant extracts.

Snails exposed to *A. indica* exhibited behaviours that suggested they had been adversely affected by these plants. The snails were weak and could neither eat nor retract into their shells. They exhibited excessive mucus secretion and cessation of feeding. Increased mucus production followed by increased mucus secretion as observed in this study, is one of the first reactions of gastropods to many stressors, including mechanical stimuli or chemical irritation caused by molluscicidal chemicals [17,18,19]. One effect of the extruded mucus is to form a protective barrier preventing direct contact between the toxin and the epithelia of the skin or digestive tract, to reduce the toxicity of the chemicals [19].

The LD_50_ of the extracts was determined to measure the dose that killed or immobilised 50% of the target organisms within the treatment period. LD_50_ figures are frequently used as a general indicator of a substance’s acute toxicity [20]. A large LD_50_ means it takes a large quantity of the material to cause a toxic response [21]. LD_50_ < 100 mg/L indicates that the substance is highly toxic, LD_50_ > 100 < 500 mg/L indicates that the substance is moderately toxic, LD_50_ > 500 < 1000 mg/L indicates that the substance is weakly toxic while LD_50_ > 1000 mg/L indicates that the substance is non-toxic [22]. Among the extracts used on adult snails, *A. indica* oil extract was highly toxic on adult snails; LD_50_< 100mg/L. *A. catechu* aqueous extract was weakly toxic; LD_50_> 100< 500mg/L while *A. catechu* ethanol extract was nontoxic on the adult snails; LD_50_>1000mg/L. When these extracts were tested on the egg masses, they were all highly toxic with *A. indica* being efficacious. For a plant to be considered a molluscicide, it should be registered in concentrations of up to 100 mg/L [23].

## Conclusion

Freshwater snails cause enormous problems in endemic areas and that inadequate control can lead to serious problems affecting inhabitants of such areas. Existing control measures are not enough to deal with emergence or outbreaks of diseases caused by freshwater snails. Therefore, continued research including using plants-based products is important to produce botanical molluscicides that are cheap, less toxic and effective to control freshwater snail population. The intermediate hosts play an essential role in the parasite life cycle. Molluscicides are therefore very crucial for controlling schistosomiasis if appropriately used. This research shows that the seed oil extract of *A. indica* had a high molluscicidal effect on *Bulinus* sp while the aqueous and ethanol extract of *A. catechu* were less toxic, and the extracts had an inhibitory effect on the development of the egg masses. For this reason, the use of plant molluscicides may be one of the veritable means of controlling schistosomiasis and other trematode infections. In addition, endemic communities are likely to accept the use of local indigenous plants since they are familiar with their properties and growth characteristics.

## Supporting Information

S1 Data. (XLSX)

## Acknowledgments

The authors thank the Head of Department of Biological Sciences laboratory for his assistance during the study.

## Author Contributions

KHT and AG, conceived the idea and designed the study. JEEE, GA, and NHN carried out the field collection and laboratory testing, and KHT performed the analysis. JEEE and GA wrote the first draft of the manuscript and was corrected by THK. All the authors contributed to the manuscript and reviewed the final manuscript for publication. All the authors read and approved the final manuscript.

